# Locating and dating land cover change events in the Renosterveld, a Critically Endangered shrubland ecosystem

**DOI:** 10.1101/2020.09.29.318568

**Authors:** Glenn R. Moncrieff

## Abstract

Land cover change is the leading cause of global biodiversity decline. New satellite platforms allow monitoring of habitats in increasingly fine detail, but most applications have been limited to forested ecosystems. I demonstrate the potential for detailed mapping and accurate dating of land cover change events in a highly biodiverse, Critically Endangered, shrubland ecosystem - the Renosterveld of South Africa. Using supervised classification of Sentinel 2 data, and subsequent manual verification with very high resolution imagery, I locate all conversion of Renosterveld to non-natural land cover between 2016 and 2020. Land cover change events are further assigned dates using high temporal frequency data from Planet labs. 478.6 hectares of Renosterveld loss was observed over this period, accounting for 0.72 % of the remaining natural vegetation in the region. 50% of change events were dated to within two weeks of their actual occurrence, and 87% to within two months. Change often preceded the planting and harvesting seasons of rainfed annual grains. These results show the potential for new satellite platforms to accurately map land cover change in non-forest ecosystems, and detect change within days of its occurrence. There is potential to use this and similar datasets to automate the process of change detection and monitor change continuously.

## Introduction

Land cover change is the leading global cause of species extinctions (Brondizio et al., 2019). Humans have significantly altered 75 % of the earth’s surface, with over one third of the terrestrial land surface currently being used for cropping or animal husbandry (FAO, 2020; Monfreda et al., 2008). Hotspots of biodiversity have on average been more altered and degraded than other areas despite their importance (Brondizio et al., 2019). The Cape Floristic Region (CFR) of South Africa is one such global biodiversity hotspot and UNESCO world heritage site (Myers et al., 2000). The CFR covers 90 000 km^2^ and is home to around 9400 plant species, 68 % of which are endemic (Manning and Goldblatt, 2012). This incredible plant species richness makes the CFR the most floristically diverse region of the world outside of the tropics. The CFR is characterised by very high fine-scale habitat partitioning among species (beta diversity) and turnover of species in space (gamma diversity) (Cowling, 1990). Species within the CFR are thus inherently vulnerable to extinction through habitat loss, as many have narrow environmental tolerances and small range sizes (Cowling and McDonald, 1998; Enquist et al., 2019).

Within the CFR, the vegetation characteristic of the lowlands is known as Renosterveld. Renosterveld typically occurs on fertile, shale-derived soils and has a significant grass component. Due to the association of Renosterveld with soils and topography suitable for agriculture, an estimated 90-94 % of it’s original extent has been completely transformed into intensively cultivated dryland crops: predominantly wheat, canola and barley (Dept of Agriculture, Western Cape, 2020; Rouget et al., 2003; Skowno et al., 2019). Sheep are grazed on fallow fields and rotational crops as well as Renosterveld patches, further degrading the few remaining intact fragments. As a result of this historic land cover change and habitat degradation the Renosterveld contains perhaps the highest concentration of threatened plant species of any continental region globally (Brummitt et al., 2015; Curtis et al., 2013; Humphreys et al., 2019; Raimondo, 2011). Within the CFR as a whole 1893 plant species are threatened with extinction, with many of these restricted to Renosterveld (Skowno et al., 2019). Of the 29 Renosterveld vegetation types described, 17 are listed as as either Critically Endangered, Endangered or Vulnerable according to the National Environmental Management: Biodiversity Act No 10 of 2004 (NEMBA). This act is intended to safeguard these ecosystems, ensuring formal protection and mandating restrictions on any loss of natural habitat. Further protection is given by the Conservation of Agricultural Resources Act, 1983 (Act No. 43 of 1983) (CARA). This act precludes the cultivation of virgin soil, defined as soil undisturbed for over 10 years, without prior authorization.

Despite this legal protection, and concerted conservation efforts (e.g the Cape Action for People and the Environment programme (CAPE) and the Overberg Renosterveld Conservation Trust), land cover transformation within the Renosterveld continues unabated (Cape Action for People and the Environment, 2009; Rouget et al., 2014). Recent estimates suggest that up to 1 % of what remains of the Renosterveld is being lost each year between 1990 and 2014, with extermination expected to be completed in less than 75 years (Skowno et al., 2019). Increasingly, the fragments of Renosterveld which do remain occur within areas that are less suitable for agriculture, with steep slopes or rocky soils (Kemper et al., 2000), thus it was thought that historical rates of vegetation loss were unlikely to be maintained. However, improving technology and deteriorating economic conditions can render areas previously thought to be of minimal agricultural value vulnerable (Kemper et al., 2000; McDowell and Moll, 1992; Rouget et al., 2003). In order to prevent the loss of the last remaining fragments of Renosterveld, conservation management agencies and environmental law enforcement require timely and accurate information on the location and timing of land cover transformation. This information can help identify the proximal and ultimate drivers of vegetation loss and identify areas that may be susceptible to future change. In recent years new algorithms and an increase in the availability of remotely sensed data have improved the temporal and spatial resolution at which land cover change can be monitored (De-Vries et al., 2015; Verbesselt et al., 2012; Zhou et al., 2019; Zhu and Woodcock, 2014). These improvements have been translated into platforms that are in use by practitioners to aid monitoring of land cover change globally (e.g. Global Forest Watch).

Continuous land cover change monitoring is typically developed for use in forest ecosystems (Hansen et al., 2013; Tang et al., 2019). These systems are well suited for this application, as land cover change involves a very large shift in vegetation - from closed canopy forest to bare ground - comparatively easy to detect in the spectral information returned from satellites. Open-canopied and low tree cover ecosystems such as grasslands and shrublands have received less attention, presumably because land cover change in these systems is harder to detect. However, these ecosystems are the dominant land cover type in sub-Saharan Africa, make up >40 % of the global total ecosystem organic carbon and contain a substantial proportion of the world’s biological diversity (Ciais et al., 2014). There is therefore an urgent need to assess the potential for continuous land cover change monitoring in non-forest ecosystems and improve the accuracy of available methods.

Here I use multi-temporal high and very-high resolution remote sensing imagery to map and accurately date land cover change events over a 4 year period from 2016 to 2020 in highly biodiverse, highly threatened, shrubland habitats within the Renosterveld of South Africa. I outline a workflow using freely available and open source tools to automatically identify areas of potential change and confirm change events. I attempt to assign a date to each land cover change event and describe the feasibility of thereof. I describe the spatial and temporal patterns of land cover change in the Renosterveld in relation to the drivers of change. The resulting dataset of located and dated land cover change events provides training data for algorithm development to accelerate the advancement of land cover change monitoring in the Renosterveld and similar ecosystems.

## Methods

Renosterveld is restricted to fertile soils in the lowlands of southern and south-western South Africa (Figure 1). This study is limited to the lowlands Renosterveld occurring within the boundaries of the Overberg district municipality, an area of 12 241 km^2^. The Overberg region was selected due to the availability of recently compiled data on the extent of remaining natural land cover, familiarity with the region and the presence of local conservation partners. Five Renosterveld vegetation types occur within the Overberg: Eastern Rûens Shale Renosterveld, Central Rûens Shale Renosterveld, Western Rûens Shale Renosterveld, Rûens Silcrete Renosterveld and Breede Shale Renosterveld (Mucina and Rutherford, 2006). All these ecosystems are listed either as Critically Endangered or Endangered under NEMBA. Together they form the natural land cover of 4711 km^2^ of the Overberg. The map of remaining natural vegetation in the Overberg is based on the combination of data on agricultural field boundaries with 2005 SPOT imagery to create a map of areas classified as transformed, degraded or natural at 1:50 000 scale (Cape Action for People and the Environment, 2009). This analysis identified 656km^2^ of the original extent of Renosterveld vegetation in the Overberg remaining as either degraded or natural. The analysis of land cover change presented here is limited to this area. A multi-step process was undertaken to 1) identify areas where Renosterveld was potentially lost between 2016 and 2020, 2) validate each site of potential loss to create an accurately digitized map of loss for this time period, and 3) for each location where loss was observed, assign a date on which this loss occurred. Figure 2 provides an overview of this method which is explained in detail below.

**Fig. 1.**
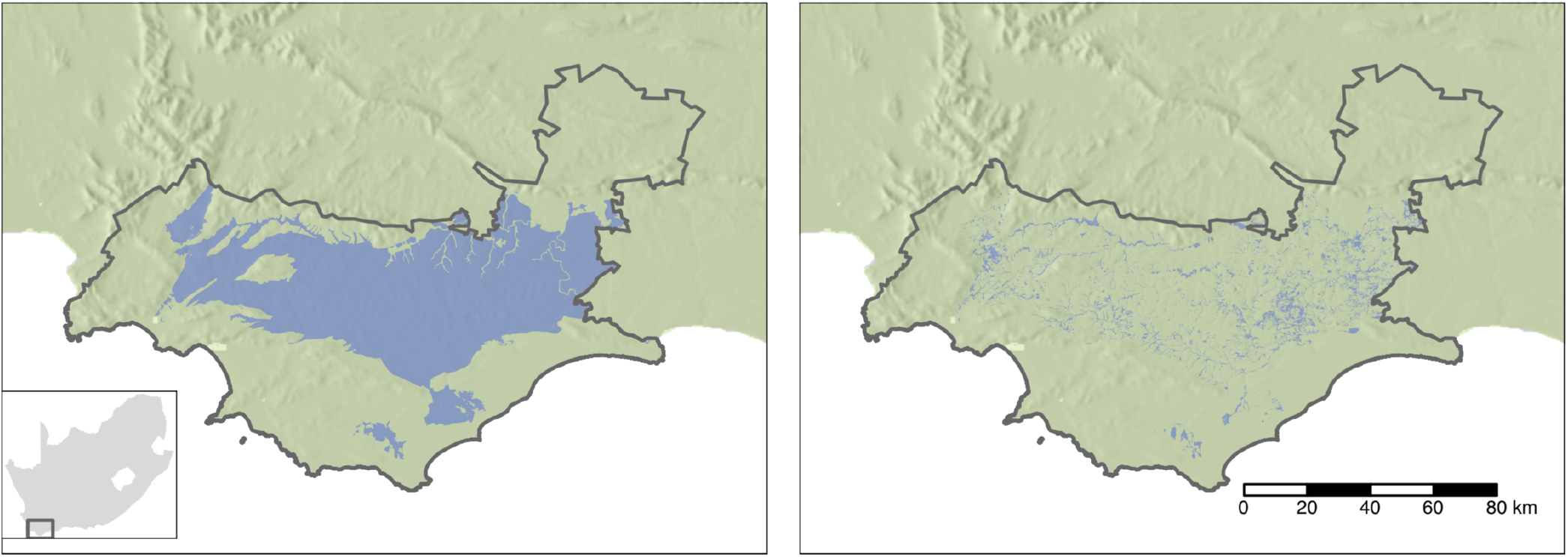
The Overberg district in South Africa. The left panel shows the natural extent of lowlands Renosterveld in the district and the right panel the remaining natural and degraded fragments

**Fig. 2.**
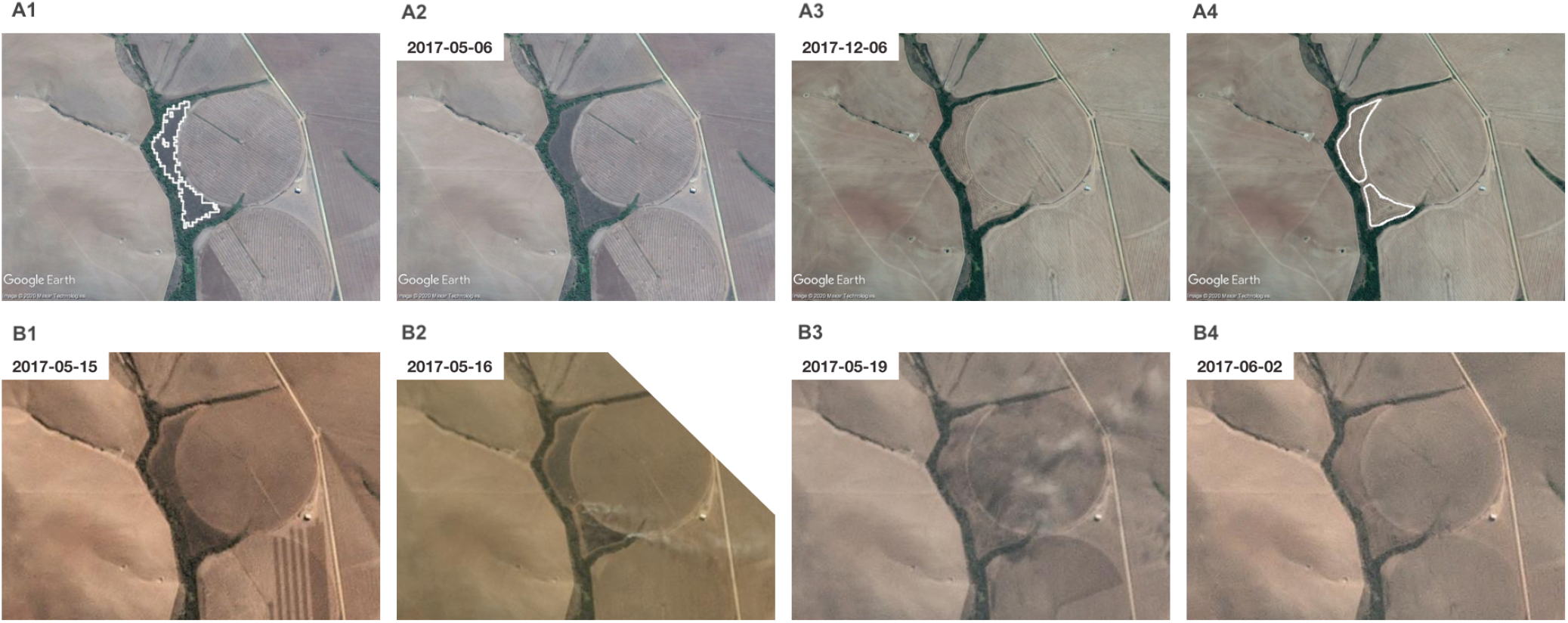
Outline of the methodology used for A) accurate mapping of Renosterveld loss and B) obtaining dates for the occurrence of change events. The output from a random forest classification outlines an area of potential Renosterveld loss (A1). Very-high resolution Google Earth imagery confirms the presence of Renosterveld on 2017-05-06 (A2) and its absence on 2017-12-06 (A3). With the help of very-high resolution imagery an accurate outline of the areas lost is obtained (A4). The time-series of PlanetScope imagery confirms 2017-05-15 as the last day that the presence of Renosterveld can be confirmed (B1). Vegetation removal is ongoing on 2017-05-16, suggested by the presence of smoke (B2). By 2017-05-19, no Renosterveld remains (B3), and this is confirmed by later imagery from 2017-06-02 (B4).

### Identifying potential change

Machine learning classification of Sentinel 2 imagery was used to identify areas of potential Renosterveld loss between 2016 and 2020, supervised using a manually created set of training data points. Sites with stable Renosterveld (no loss between February 2016 and January 2020), stable agriculture (cultivated by February 2016) and Renosterveld loss (loss of Renosterveld between February 2016 and January 2020) were identified using very high resolution Google Earth imagery and 50cm resolution aerial photography (NGI, 2020). Any area under cultivation by February 2016 was considered stable agriculture, as it is assumed impossible for agriculture to revert to natural land cover over this time frame (Heelemann et al., 2012, 2013; Ruwanza, 2020). Points were chosen to cover a variety of agricultural land uses types using a regional map of crop types (Western Cape Department of Agriculture, 2013). Points for stable Renosterveld were located in areas with apparently intact Renosterveld in January 2020. It is impossible to confirm that sites were pristine, primary Renosterveld from remotely sensed imagery alone. However, the presence of shrubs and heterogeneous plant composition characteristic of Renosterveld stands in stark contrast to the monocultures or bare soil characteristic of agricultural land, and can thus be discerned with confidence by a trained eye. The reliability of Renosterveld identification from aerial imagery alone was confirmed by multiple ad hoc excursions. Locations of loss of Renosterveld between February 2016 and January 2020 were created from identifying areas where the presence of Renosterveld could be confirmed in February 2016 and the presence of non-natural land cover could be confirmed in January 2020. The location of sites where change was suspected to have occurred between these dates was provided by local conservation partners. The final dataset of labelled sites consists of 2177 sites for stable Renosterveld, 2336 sites for stable agriculture, and 909 site of Renosterveld loss. QGIS 3.10 and Google Earth were used for image interpretation and digitization.

Input features were derived from Sentinel 2 L1C data from 2016-02-06 and 2020-01-11. The entire study region falls within a single Sentinel 2 track (sensing orbit number 121) and thus images from a single date could be used for the beginning and end of the monitoring period, ensuring relatively homogeneous atmospheric conditions over the region. Dates were selected based on the availability of favourable atmospheric conditions and the absence of clouds, while mid-to late-summer was preferred due to the strong visual contract between fallow fields and Renosterveld at this time of year. Three spectral indices were calculated for each image: normalised difference vegetation index (NDVI), normalised difference red-edge index (NDRE), and the enhanced vegetation index (EVI). In addition to these, a single measure of spectral distance between these two images for each pixel was calculated using the spectral angle mapper algorithm (Yuhas et al., 1992). Combining this with the calculated VI’s and bands 2,3,4,5,6,7,8,8A, 11 and 12 from each image results in a total of 27 input features. These features along with the labelled points were used to train a Random Forest classifier. The labelled data were split into 5 folds based on location, and spatial k-fold cross validation was used to assess performance (Ploton et al., 2020). The trained model was then used for prediction over the entire study region. The output of this prediction was filtered to retain only predictions of Renosterveld loss and masked to areas mapped as degraded or natural in 2008. Only areas of change larger than 0.1 ha or roughly ten contiguous pixels were retained for subsequent investigation. Google Earth Engine was used for data preparation, model fitting and evaluation (Gorelick et al., 2017).

### Precisely locating change

Each of these potential land cover change events was then manually validated using very-high resolution Google Earth imagery, 50cm resolution aerial photography (NGI, 2020) and high resolution (3-5m) PlanetScope imagery from Planet labs (PlanetTeam, 2019). Every contiguous output parcel was checked to ensure that 1) land-cover was indeed non-natural / agriculture at the end of the monitoring period, 2) intact Renosterveld was present at the start of the monitoring period, and 3) land was uncultivated at least 10 years prior to the start of the monitoring period and 4) that the change took place between February 2016 and January 2020. If these four criteria were met, the accuracy of the output outline was improved by manually digitising by the area of Renosterveld lost at 1:50 000 scale. Each of these output Renosterveld change events was reviewed and confirmed by an second independent assessor and a local vegetation expert.

### Dating of change events

High temporal frequency PlanetScope imagery was then used to determine the date on which loss occurred for each parcel where loss was confirmed between February 2016 and January 2020. The removal of Renosterveld within a parcel can take multiple days or weeks to be completed, and even if removal occurs instantaneously multiple days or weeks can elapse before a clear image becomes available in which it is possible to confidently confirm that natural vegetation is no longer present. Two dates were therefore assigned to each parcel to describe the temporal nature of Renosterveld loss: the latest date on which it was possible to confirm that intact Renosterveld was still present, and the earliest date on which it was possible to confirm that intact Renosterveld was no longer present. Determining these dates depends on the revisitation frequency of satellites platforms, atmospheric conditions which determine the image useability, and the nature of land cover change (Figure 2). The near-daily global imaging capability of the Planet satellite constellation and low cloud and haze over this region for much of the year provide sufficient images for these dates to be determined within a few days of each other for many of the Renosterveld loss events detected. The mechanism through which Renosterveld is converted to non-natural land cover, and whether change is continuous or punctuated, influences how discernible land cover change is when examining satellite image time-series’. This in turn influences how accurately loss events can be dated. Given an abrupt disturbance that removes a large amount of aboveground biomass and disturbs the soil surface - such as ploughing - the last date of Renosterveld presence and first date of it’s absence can be determined within a few days if sufficient imagery is available. Gradual vegetation degradation through continual overgrazing can results in the loss of Renosterveld over a period of multiple years or even decades. This type of land cover change is poorly captured by the approach used here. To date each parcel of Renosterveld loss, the approximate month and year in which loss occurred was determined using monthly global surface reflectance basemaps available from Planet labs. Once this month was determined, time series’ of daily imagery were examined to determine the last date of Renosterveld presence and first date of it’s absence (Figure 2) using the Planet Explorer online platform. Only those Renosterveld loss parcels in which these two dates could be confidently determined within 10 weeks of each other were used in subsequent exploration and analysis of temporal patterns of land cover change.

## Results

The random forest classification predicts an area of 3644 ha of potential Renosterveld loss between February 2016 and January 2020. The fitted model achieves an overall accuracy of 90 %. For the Renosterveld loss, stable Renosterveld and stable agriculture classes producers accuracy was 72%, 94% and 95%, and users accuracy was 81%, 92% and 91% respectively. Despite the accuracy of this algorithm, many instances of false positives of Renosterveld loss are observed when manually comparing model output to very high resolution satellite imagery. Fewer instances of false negatives occurred. Common causes of false identification of Renosterveld loss were wildfires within stable Renosterveld and stable agriculture miss-classified as Renosterveld loss.

After manual removal of false detections and accurate digitisation of boundaries, the final dataset of confirmed Renosterveld loss between February 2016 and January 2020 contained 268 contiguous events covering 478.6 ha. 87 % of the area identified as Renosterveld loss by the random forest was not confirmed and hence does not appear in the final dataset. While the majority of these locations were indeed false positives, others suggest potential change. However, if it was not possible to confirm that Renosterveld removal or degradation resulted in vegetation transitioning from intact to completely lost between February 2016 and January 2020, the site was not included in the final dataset. This criterion means that the a higher percentage of the area identified by the random forests is indeed Renosterveld loss, and that the true area lost is higher than the confirmed 478.6 ha.

The confirmed loss of 478.6 ha represents 0.73 % of the remaining 65 684 ha of Renosterveld mapped as either degraded or natural (Cape Action for People and the Environment, 2009). It is likely that this is a significant overestimate of the extent of intact Renosterveld extent in 2016 and thus the true proportion lost is slightly higher than this figure. Most of the events that result in Renosterveld loss are relatively small, with 57% smaller than 1 ha and 93% smaller than 5 ha (Figure 3). A few large events (18) greater than 5 ha account for 41% of the total area lost.

**Fig. 3.**
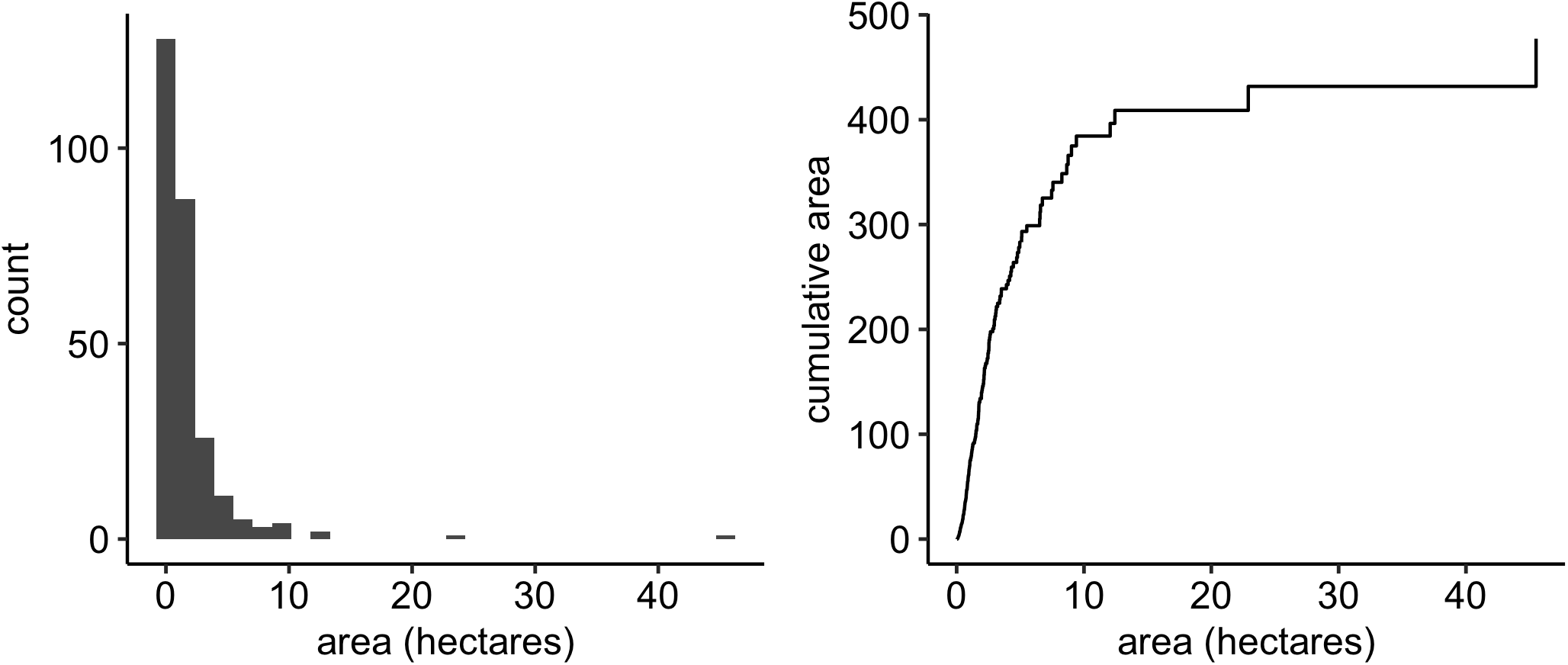
Histogram of the size of Renosterveld loss events (left), and the cumulative area lost by events of increasing size (right).

The date of the 232 (87%) confirmed Renosterveld loss events was determined to within 60 days (the time elapsed between the final confirmed date of Renosterveld presence and first confirmed date of Renosterveld absence) - figure 4. Of these events the date of a further 134 (50%) could be determined to within 14 days, and 55 (21%) within 7 days. Figure 5 shows the geographic distribution of Renosterveld loss events along with their size and median date of the loss event (the date halfway between the final confirmed date of Renosterveld presence and first confirmed date of Renosterveld absence). In general, Renosterveld loss occurred earlier in the western Overberg than the eastern Overberg during this time period. While spread throughout the region, clusters of land cover change are evident in multiple regions (from west to east): northwest of the town of Botrivier, north of the Caledonberg between the town of Caledon and Helderstroom, the Karingmelksrivier west of Napier, southeast of Greyton, east of Protem and both the eastern and western sides of the Breede river north of Malgas.

**Fig. 4.**
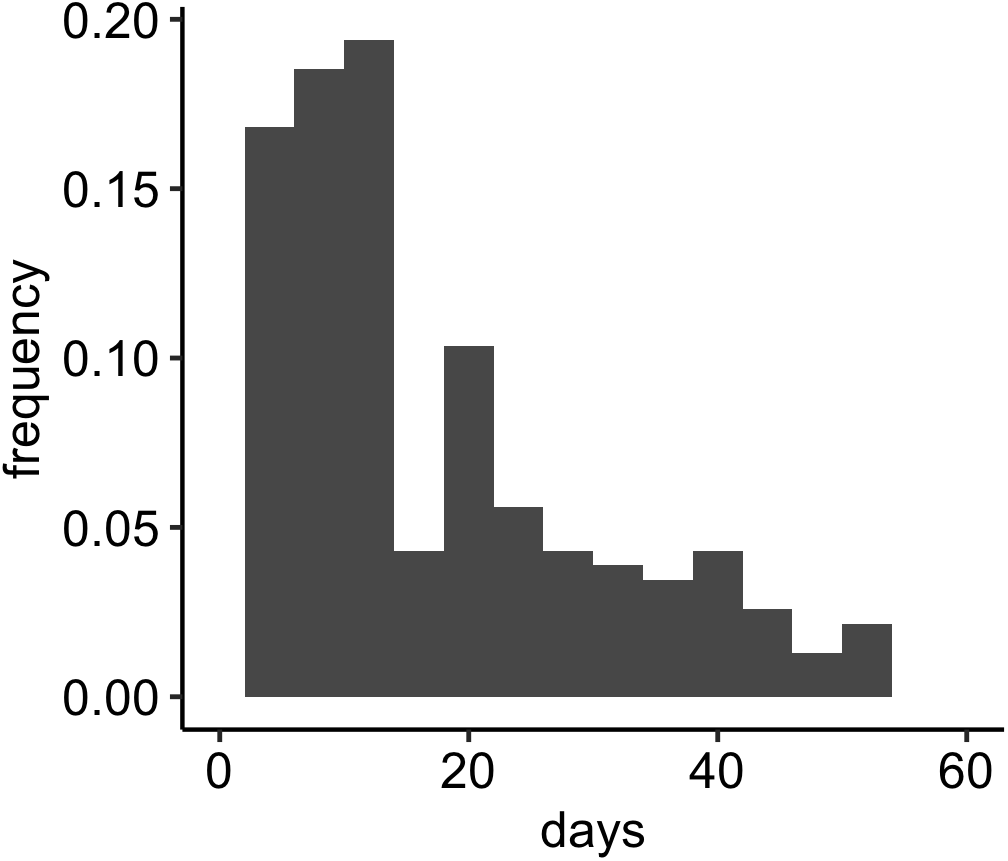
Frequency histogram of days elapsed between the last days of confirmed Renosterveld presence and the first day of confirmed absence

**Fig. 5.**
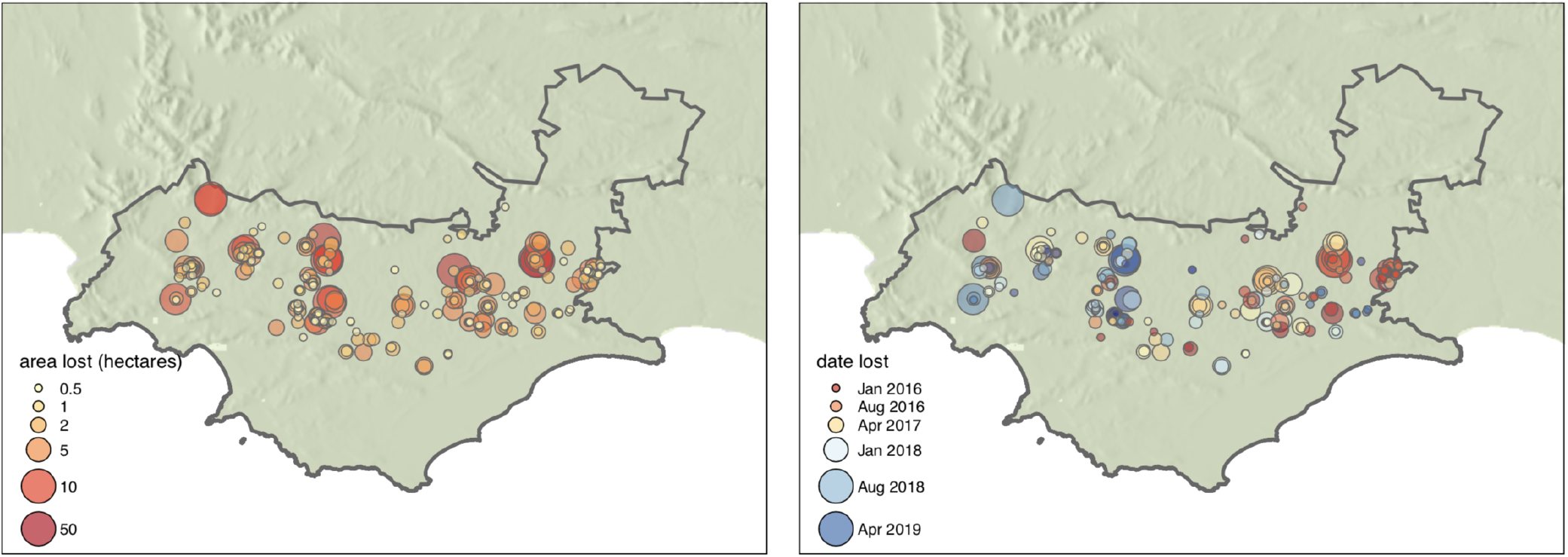
The spatial distribution of Renosterveld loss observed in the Overberg district between February 2016 and Januray 2020. Point colour and size indicate the size of loss events (left), and the date of loss and event size respectively (right).

Renosterveld loss events occurred throughout the year, though across all year the months with the highest concentration of events and the most area lost were February - March, and August to September (Figure 6). Little land cover change took place in the months of December-January and July. Between 2016 and 2020, 2017 saw the most loss events (despite a large area lost in 2016 due to a few related, large events). No Renosterveld loss was detected and dated in the latter half of 2019.

**Fig. 6.**
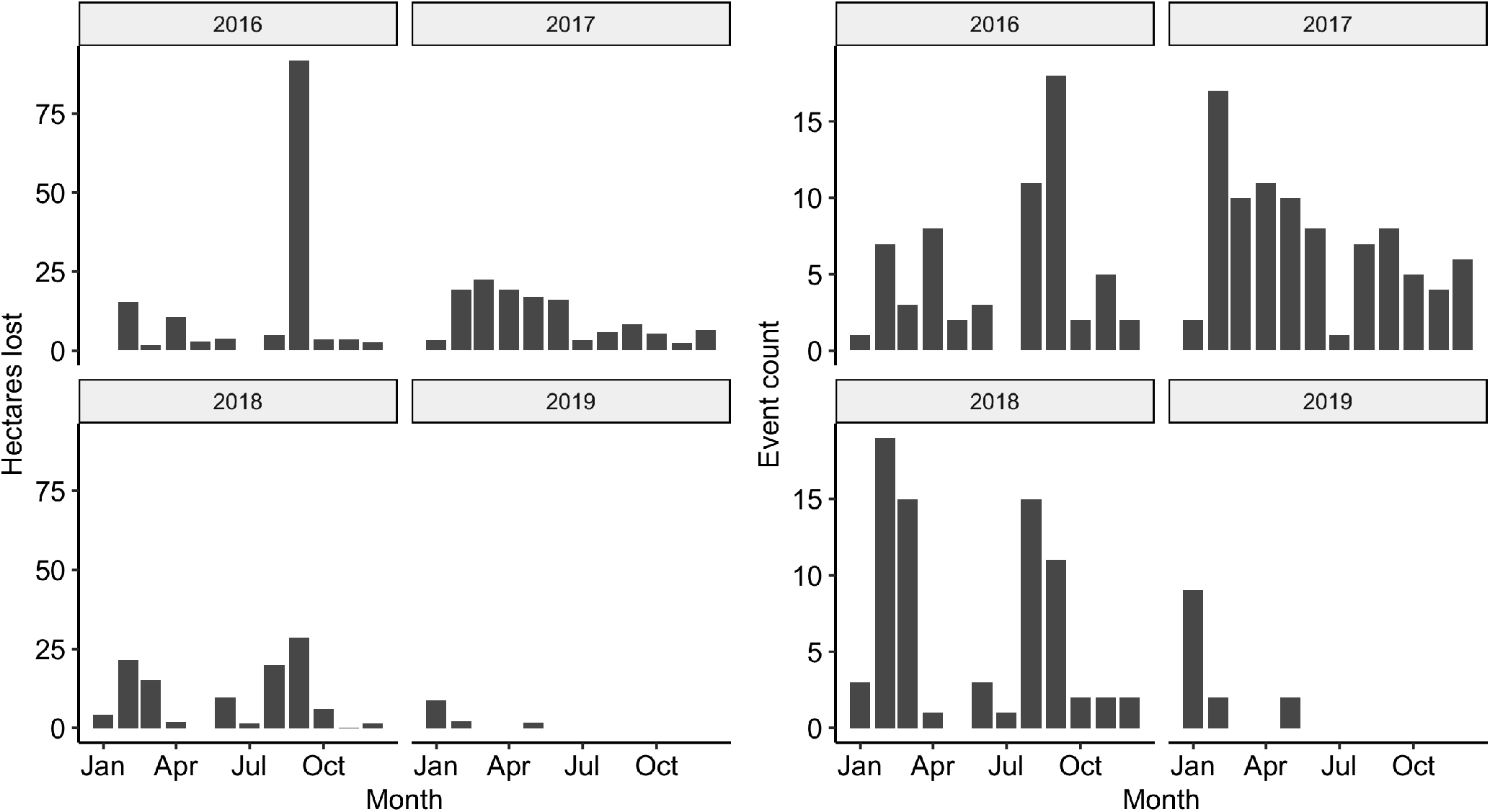
Hectares of Renosterveld lost (left) and number of loss events (right), defined as spatially contiguous polygons, by month and year over the study period.

## Discussion

The vegetation monitored in this study is of exceptionally high conservation value. The density of species threatened with extinction is almost unmatched globally, and every remnant of natural habitat is of enormous value in ensuring the continued survival of the flora and fauna of the Renosterveld (Kemper et al., 1999, 2000; Skowno et al., 2019; Von Hase et al., 2003). The loss of 478 hectares of intact Renosterveld between February 2016 and January 2020 shows that the current approach to conservation in the region is failing in its goal of preventing further loss of what remains of the Renosterveld. The ability to monitor the daily progression of land cover change in this region demonstrated here provides information with which to better understand the drivers of Renosterveld loss and the potential for improved enforcement of existing protections. Large inter- and intra-annual variability in rates of loss suggests that actors responsible for land cover change are doing so in response to external factors. Accurate dating of individual events aids in diagnosing these factors and potential future mitigation (Qin et al., 2019). For law enforcement and others involved in efforts to prevent further loss, the ability to continuously monitor ongoing change will serve as a strong deterrent, and aid in the deployment of existing resources.

Until recently frequent monitoring of land cover change at high (<30m) spatial resolution has either been impossible or prohibitively expensive. This has precluded the detailed study of land cover change dynamics in highly fragmented and non-forest ecosystems. With the growing time series of data freely available from Sentinel 2 and the unprecedented temporal resolution of the Planet labs Dove constellation it has become possible to reconstruct accurate timelines of land cover change and continuously monitoring ongoing change in many of the world ecosystems (Grabska et al., 2020; Zhou et al., 2019). The approach applied here produced a detailed description of the patterns of land cover change in space and time, allowing 50% of events to be dated within 2 weeks of their occurrence, and the loss of fragments as small as 300 m^2^ to be detected in a highly fragmented, low-biomass shrubland ecosystem. A significant amount of expert knowledge and manual image interpretation is required to locate and date land cover change events in this ecosystem. However, screening of areas in which change is likely through the rapid creation of a simple training dataset and application of standard classification algorithms greatly improved the efficiency of this task.

This map of Renosterveld change events with associated dates provides a unique dataset with the potential to automate monitoring in the region. Existing approaches to continuous monitoring and detection of change almost exclusively apply an unsupervised approach to detecting land cover change (e.g. DeVries et al. (2015); Verbesselt et al. (2012); Zhou et al. (2019); Zhu and Woodcock (2014)). Abrupt changes in time-series’ of vegetation activity are interpreted as land cover change in this framework. Natural processes that may cause surface reflectance to change, such as natural wildfire or treefall, will be missclassified. Using a dataset in which only land cover change events caused by anthropogenic processes are included to train an algorithm in a supervised manner may separate the signal of unnatural disturbance from that of natural disturbance. The additional knowledge of the date of each event enables training algorithms that will be robust to seasonal changes in vegetation, and hence can be applied continuously as new data becomes available rather than annually (Kennedy et al., 2010, 2014). Recent advances in the supervised classification of land cover from annual time-series of Sentinel 2 data provide an excellent starting point for experimentation (Pelletier et al., 2019; Rußwurm and Körner, 2019)

All the vegetation types monitored in this study are listed as Critically Endangered or Endangered ecosystems in term of the South Africa’s NEMBA act. The act stipulates that removal of vegetation in these ecosystems requires prior authorization and a basic environmental assessment. Very few such authorizations were granted in this region over the period studied and therefore it is likely that most of the change reported here can be considered unlawful. Furthermore, it was confirmed that changes affected virgin soil that had remained undisturbed for at least 10 years (but in most cases far longer) as per the CARA act. This precludes the invocation of the common defense that old fields rather than intact natural vegetation are being targeted. The information obtained here is being used to aid ongoing investigations and has assisted in the application of administrative penalties for NEMBA contraventions. However, as demonstrated by the rate of change reported here, Renosterveld loss is ongoing. Corrective action to restore lost habitat cannot compensate for the loss of primary natural vegetation to agriculture in this ecosystem (Heelemann et al., 2012, 2013; Ruwanza, 2020). The only effective mitigation measures are intervention while habitat destruction is ongoing or the deterrence of future damage. The implementation of real-time monitoring as described above could allow officials to detect habitat loss soon enough after its initiation that intervention may occur before vegetation removal is completed. Even if this is not possible, reducing the time between the occurrence of an event and expert ground-truthing would aid in the collection of evidence for use in prosecution. Curbing the Renosterveld loss resulting from the accumulation of multiple small events may require a different strategy than the 41% that is lost to a few actors removing areas greater than 5 ha in a few incidences. The realization that remote sensing technology is in use to monitor threatened ecosystems may itself serve as a deterrent of potential infringements (Kelly and Kelly, 2014).

The nature of Renosterveld loss observed in this study was limited to acute vegetation change within a period of a few days to a few years. Though it is not possible to diagnose exactly the cause of change for each event, the principle factor was ploughing for cultivation. Only a few instance of overgrazing combined with fire were observed to completely transform Renosterveld in this timeframe. The intra-annual patterns of Renosterveld loss reflect patterns of agricultural activity in the region, with the peaks of loss occurring in the months prior to the planting of grains in April and May and before harvesting in October and November. Inter-annual variability may be related to high variability in yields in this region of predominantly dryland agriculture (GrainSA, 2020). Poor rainfall over this region over certain years in the study period (2017 and 2019) and macroeconomic factors such as fluctuationd in land value may explain some of the patterns seen over the study period, but a longer time series of change would be required to examine these drivers in detail (Skowno et al., 2019; Sousa et al., 2018).

The efficiency of the painstaking work of accurately digitizing areas of potential loss and dating individual change events could be improved through the application of improved algorithms and a larger, more diverse training data set (Pelletier et al., 2017, 2019; Rußwurm and Körner, 2019). But image interpretation will continue to require the expertise of an individual with familiarly of the ecosystem under consideration. Land cover change that occurs slowly over decades or longer as a result of long-term degradation through inappropriate fire regimes, overgrazing or invasive alien species is not captured on the time scale studied here. These factors contribute significantly to the long-term decline of Renosterveld (Topp and Loos, 2019). This may account for the discrepancy between the annual rate of Renosterveld loss reported here of 0.18 % per annum and the annual rate reported by Skowno et al. (2019) of approximately 1%. This limitation may be overcome as longer time series’ of data from Sentinel 2 and Planet labs become available, though it will remain impossible to assign a precise date of change to these type of events. In order to slow and ultimately halt the ongoing loss of Renosterveld a co-ordinated approach is needed. The process described here can provide helpful information to understand the drivers of loss and a lead to the development of tools to monitor change in real-time. This information is already being used by environmental enforcement authorities to aid investigations and deter future infringements. But progress will be limited unless a systematic approach addressing the underlying causes is developed and behavioural change is incentivised. This has been attempted before (Von Hase et al., 2003), and while these efforts have borne fruit, they have not proved sufficient. Efforts to mobilize additional funds and test new approaches to conservation in the region - including using the methods described here - are currently underway.

## Conclusions

The Overberg Renosterveld has been identified as South Africa’s most threatened ecosystem. The rate of loss of 478 ha over 4 years observed here is unsustainable and leading to the extinction of this vegetation type. Accurate mapping and reconstruction of the timeline of events surrounding land cover change in the region is possible using remotely sensed data. These data can be used to aid law enforcement operations, monitor ongoing change and develop plans to mitigate future loss. This demonstrates the possibility of similar data being collected for other shrublands and low tree cover vegetation, and contributing to improved management of threatened ecosystems globally.

## ACKNOWLEDGEMENTS

I am very grateful for the support and encouragement given by Odette Curtis and her passion for conservation in this region. Keletso Moilwe and Lungile Khuzwayo assisted with digitization. Multiple individuals from the Western Cape Department of Environmental Affairs and Development Planning are acknowledged for their input, in particular, those from the Environmental Law Enforcement directorate. This research was supported by the National Research Foundation of South Africa through (Grant No. 118593) as part of the RReTool: Rapid and repeatable tools for monitoring and mitigating global change impacts on natural resources project. The funders had no role in study design, data collection and analysis, decision to publish, or preparation of the manuscript.

## DATA AVAILABILITY

Code and data used in this analysis is available at www.github.com/GMoncrieff/renosterveld-change. Due to ongoing investigations location data has been removed to protect the identity of landowners where necessary.

## Notes

https://github.com/GMoncrieff/renosterveld-change

### Competing Interest Statement

The authors have declared no competing interest.

## Bibliography

E. S. Brondizio, J. Settele, S. Díaz, and H. T. Ngo. Global assessment report on biodiversity and ecosystem services of the Intergovernmental Science-Policy Platform on Biodiversity and Ecosystem Services. IPBES Secretariat: Bonn, Germany, 2019.

N. A. Brummitt, S. P. Bachman, J. Griffiths-Lee, M. Lutz, J. F. Moat, A. Farjon, J. S. Donaldson, C. Hilton-Taylor, T. R. Meagher, S. Albuquerque, E. Aletrari, A. K. Andrews, G. Atchison, E. Baloch, B. Barlozzini, A. Brunazzi, J. Carretero, M. Celesti, H. Chadburn, E. Cianfoni, C. Cockel, V. Coldwell, B. Concetti, S. Contu, V. Crook, P. Dyson, L. Gardiner, N. Ghanim, H. Greene, A. Groom, R. Harker, D. Hopkins, S. Khela, P. Lakeman-Fraser, H. Lindon, H. Lockwood, C. Loftus, D. Lombrici, L. Lopez-Poveda, J. Lyon, P. Malcolm-Tompkins, K. McGregor, L. Moreno, L. Murray, K. Nazar, E. Power, M. Q. Tuijtelaars, R. Salter, R. Segrott, H. Thacker, L. J. Thomas, S. Tingvoll, G. Watkinson, K. Wojtaszekova, and E. M. N. Lughadha. Green Plants in the Red: A Baseline Global Assessment for the IUCN Sampled Red List Index for Plants. PLOS ONE, 10(8):e0135152, Aug. 2015.

Cape Action for People and the Environment. Overberg District Transformation Layer 2009 - Biodiversity BGIS, 2009. URL https://bgis.sanbi.org/SpatialDataset/Detail/385.

P. Ciais, C. Sabine, G. Bala, L. Bopp, V. Brovkin, J. Canadell, A. Chhabra, R. DeFries, J. Galloway, and M. Heimann. Carbon and other biogeochemical cycles. In Climate change 2013: the physical science basis. Contribution of Working Group I to the Fifth Assessment Report of the Intergovernmental Panel on Climate Change, pages 465–570. Cambridge University Press, 2014.

R. M. Cowling. Diversity Components in a Species-Rich Area of the Cape Floristic Region. Journal of Vegetation Science, 1(5):699–710, 1990.

R. M. Cowling and D. J. McDonald. Local Endemism and Plant Conservation in the Cape Floristic Region. In P. W. Rundel, G. Montenegro, and F. M. Jaksic, editors, Landscape Disturbance and Biodiversity in Mediterranean-Type Ecosystems, Ecological Studies, pages 171–188. Springer, Berlin, Heidelberg, 1998.

O. E. Curtis, C. H. Stirton, and A. M. Muasya. A conservation and floristic assessment of poorly known species rich quartz–silcrete outcrops within Rûens Shale Renosterveld (Overberg, Western Cape), with taxonomic descriptions of five new species. South African Journal of Botany, 87:99–111, July 2013.

Dept of Agriculture, Western Cape. Western Cape AgriStats, 2020. URL http://www.elsenburg.com/gis/apps/agristats/.

B. DeVries, J. Verbesselt, L. Kooistra, and M. Herold. Robust monitoring of small-scale forest disturbances in a tropical montane forest using Landsat time series. Remote Sensing of Environment, 161:107–121, 2015.

B. J. Enquist, X. Feng, B. Boyle, B. Maitner, E. A. Newman, P. M. Jørgensen, P. R. Roehrdanz, B. M. Thiers, J. R. Burger, R. T. Corlett, T. L. P. Couvreur, G. Dauby, J. C. Donoghue, W. Foden, J. C. Lovett, P. A. Marquet, C. Merow, G. Midgley, N. Morueta-Holme, D. M. Neves, A. T. Oliveira-Filho, N. J. B. Kraft, D. S. Park, R. K. Peet, M. Pillet, J. M. Serra-Diaz, B. Sandel, M. Schildhauer, I. Šímová, C. Violle, J. J. Wieringa, S. K. Wiser, L. Hannah, J.-C. Svenning, and B. J. McGill. The commonness of rarity: Global and future distribution of rarity across land plants. Science Advances, 5(11):eaaz0414, Nov. 2019.

FAO, 2020. URL http://faostat.fao.org/static/syb/syb_5000.pdf.

N. Gorelick, M. Hancher, M. Dixon, S. Ilyushchenko, D. Thau, and R. Moore. Google Earth Engine: Planetary-scale geospatial analysis for everyone. Remote sensing of Environment, 202:18–27, 2017.

E. Grabska, P. Hawrylo, and J. Socha. Continuous Detection of Small-Scale Changes in Scots Pine Dominated Stands Using Dense Sentinel-2 Time Series. Remote Sensing, 12(8):1298, Jan. 2020.

GrainSA. Crop Estimates, 2020. URL https://www.grainsa.co.za/pages/industry-reports/crop-estimates.

M. C. Hansen, P. V. Potapov, R. Moore, M. Hancher, S. A. Turubanova, A. Tyukavina, D. Thau, S. V. Stehman, S. J. Goetz, and T. R. Loveland. High-resolution global maps of 21st-century forest cover change. science, 342(6160):850–853, 2013.

S. Heelemann, C. B. Krug, K. J. Esler, C. Reisch, and P. Poschlod. Pioneers and Perches—Promising Restoration Methods for Degraded Renosterveld Habitats? Restoration Ecology, 20(1):18–23, 2012.

S. Heelemann, C. B. Krug, K. J. Esler, C. Reisch, and P. Poschlod. Soil seed banks of remnant and degraded Swartland Shale Renosterveld. Applied Vegetation Science, 16(4):585–597, 2013.

A. M. Humphreys, R. Govaerts, S. Z. Ficinski, E. Nic Lughadha, and M. S. Vorontsova. Global dataset shows geography and life form predict modern plant extinction and rediscovery. Nature Ecology & Evolution, 3(7):1043–1047, July 2019.

A. B. Kelly and N. M. Kelly. Validating the remotely sensed geography of crime: a review of emerging issues. Remote Sensing, 6(12):12723–12751, 2014.

J. Kemper, R. M. Cowling, and D. M. Richardson. Fragmentation of south african renosterveld shrublands: effects on plant community structure and conservation implications. Biological Conservation, 90(2):103–111, 1999.

J. Kemper, R. M. Cowling, D. M. Richardson, G. G. Forsyth, and D. H. McKelly. Landscape fragmentation in South Coast Renosterveld, South Africa, in relation to rainfall and topography. Austral Ecology, 25(2):179–186, 2000.

R. E. Kennedy, Z. Yang, and W. B. Cohen. Detecting trends inforest disturbance and recovery using yearly Landsat time series: 1. LandTrendr – Temporal segmentation algorithms. Remote Sensing of Environment, 114(12):2897–2910, Dec. 2010.

R. E. Kennedy, S. Andréfouët, W. B. Cohen, C. Gómez, P. Griffiths, M. Hais, S. P. Healey, E. H. Helmer, P. Hostert, M. B. Lyons, and others. Bringing an ecological view of change to Landsat-based remote sensing. Frontiers in Ecology and the Environment, 12(6):339–346, 2014.

J. Manning and P Goldblatt. Plants of the Greater Cape Floristic Region. 1: The Core Cape flora. South African National Biodiversity Institute, 2012.

C. McDowell and E. Moll. The influence of agriculture on the decline of west coast renosterveld, south-western Cape, South Africa. Journal of Environmental Management, 35(3):173–192, July 1992.

C. Monfreda, N. Ramankutty, and J. A. Foley. Farming the planet: 2. Geographic distribution of crop areas, yields, physiological types, and net primary production in the year 2000. Global Biogeochemical Cycles, 22(1), 2008.

L. Mucina and M. C. Rutherford, editors. The vegetation of South Africa, Lesotho and Swaziland. Strelitzia 19. South African National Biodiversity Institute, Pretoria., 2006.

N. Myers, R. A. Mittermeier, C. G. Mittermeier, G. A. B. da Fonseca, and J. Kent. Biodiversity hotspots for conservation priorities. Nature, 403(6772):853–858, Feb. 2000.

NGI. National Geo-spatial Information, Department of Rural Development and Land Reform, 2020. URL http://www.ngi.gov.za/index.php/what-we-do/aerial-photography-and-imagery/35-colour-digital-aerial-imagery-at-0-5m-gsd-2008-2016-and-0-25m-gsd-2017-cur

C. Pelletier, S. Valero, J. Inglada, N. Champion, C. Marais Sicre, and G. Dedieu. Effect of Training Class Label Noise on Classification Performances for Land Cover Mapping with Satellite Image Time Series. Remote Sensing, 9(2):173, Feb. 2017.

C. Pelletier, G. I. Webb, and F. Petitjean. Temporal Convolutional Neural Network for the Classification of Satellite Image Time Series. Remote Sensing, 11(5):523, Jan. 2019.

PlanetTeam. Planet Application Program Interface: In Space for Life on Earth. San Francisco, CA https://api.planet.com, 2019.

P. Ploton, F. Mortier, M. Réjou-Méchain, N. Barbier, N. Picard, V. Rossi, C. Dormann, G. Cornu, G. Viennois, N. Bayol, A. Lyapustin, S. Gourlet-Fleury, and R. Pélissier. Spatial validation reveals poor predictive performance of large-scale ecological mapping models. Nature Communications, 11(1):4540, Sept. 2020.

Y. Qin, X. Xiao, J. Dong, Y. Zhang, X. Wu, Y. Shimabukuro, E. Arai, C. Biradar, J. Wang, Z. Zou, F. Liu, Z. Shi, R. Doughty, and B. Moore. Improved estimates of forest cover and loss in the Brazilian Amazon in 2000–2017. Nature Sustainability, 2(8):764–772, Aug. 2019.

D. Raimondo. The Red List of South African plants: a global first. South African Journal of Science, 107(3-4):01–02, Apr. 2011.

M. Rouget, D. M. Richardson, R. M. Cowling, J. W. Lloyd, and A. T. Lombard. Current patterns of habitat transformation and future threats to biodiversity in terrestrial ecosystems of the Cape Floristic Region, South Africa. Biological Conservation, 112(1):63–85, 2003.

M. Rouget, M. Barnett, R. M. Cowling, T. Cumming, F. Daniels, M. T. Hoffman, and T. Rebelo. Conserving the Cape Floristic Region. Fynbos: Ecology, Evolution and Conservation of a Megadiverse Region, pages 321–336, 2014.

S. Ruwanza. Topsoil Transfer from Natural Renosterveld to Degraded Old Fields Facilitates Native Vegetation Recovery. Sustainability, 12(9):3833, Jan. 2020.

M. Rußwurm and M. Körner. Self-attention for raw optical satellite time series classification. arXiv preprint arXiv:1910.10536, 2019.

A. J. Skowno, C. J. Poole, and D. C. Raimondo. National biodiversity assessment 2018: the status of South Africa’s ecosystems and biodiversity. Synthesis Report. South African National Biodiversity Institute, Pretoria, 2019.

P. M. Sousa, R. C. Blamey, C. J. C. Reason, A. M. Ramos, and R. M. Trigo. The ‘Day Zero’ Cape Town drought and the poleward migration of moisture corridors. Environmental Research Letters, 13(12):124025, Dec. 2018.

X. Tang, E. L. Bullock, P. Olofsson, S. Estel, and C. E. Woodcock. Near real-time monitoring of tropical forest disturbance: New algorithms and assessment framework. Remote Sensing of Environment, 224:202–218, Apr. 2019.

E. N. Topp and J. Loos. Fragmented Landscape, Fragmented Knowledge: A Synthesis of Renosterveld Ecology and Conservation. Environmental Conservation, 46(2):171–179, June 2019.

J. Verbesselt, A. Zeileis, and M. Herold. Near real-time disturbance detection using satellite image time series. Remote Sensing of Environment, 123:98–108, 2012.

A. Von Hase, M. Rouget, K. Maze, and N. Helme. A fine-scale conservation plan for Cape lowlands renosterveld: technical report, 2003.

Western Cape Department of Agriculture. Crop Census, 2013.

R. H. Yuhas, A. F. Goetz, and J. W. Boardman. Discrimination among semi-arid landscape endmembers using the spectral angle mapper (SAM) algorithm. JPL, Summaries of the Third Annual JPL Airborne Geoscience Workshop. Volume 1: AVIRIS Workshop, 1992.

Q. Zhou, J. Rover, J. Brown, B. Worstell, D. Howard, Z. Wu, A. L. Gallant, B. Rundquist, and M. Burke. Monitoring Landscape Dynamics in Central U.S. Grasslands with Harmonized Landsat-8 and Sentinel-2 Time Series Data. Remote Sensing, 11(3):328, Jan. 2019.

Z. Zhu and C. E. Woodcock. Continuous change detection and classification of land cover using all available Landsat data. Remote Sensing of Environment, 144:152–171, Mar. 2014.

